# The physiological butyrylcholinesterase tetramer is a dimer of dimers stabilized by a superhelical assembly

**DOI:** 10.1101/431288

**Authors:** Miguel Ricardo Leung, Laura S. van Bezouwen, Lawrence M. Schopfer, Joel L. Sussman, Israel Silman, Oksana Lockridge, Tzviya Zeev-Ben-Mordehai

## Abstract

The quaternary structures of the cholinesterases, acetylcholinesterase (AChE) and butyrylcholinesterase (BChE), are essential for their localisation and function. Of practical importance, BChE is a promising therapeutic candidate for intoxication by organophosphate nerve agents and insecticides, and for detoxification of addictive substances. Efficacy of the recombinant enzyme hinges on its having a long circulatory half-life; this, in turn, depends strongly on its ability to tetramerize. Here, we used cryo-electron microscopy (cryo-EM) to determine the structure of the highly glycosylated native BChE tetramer purified from human plasma at 5.7 Å. Our structure reveals that the BChE tetramer is organised as a staggered dimer of dimers. Tetramerization is mediated by assembly of the C-terminal tryptophan amphiphilic tetramerization (WAT) helices from each subunit as a superhelical assembly around a central anti-parallel polyproline II helix (PRAD). The catalytic domains within a dimer are asymmetrically linked to the WAT/PRAD. In the resulting arrangement, the tetramerization domain is largely shielded by the catalytic domains, which may contribute to the stability of the HuBChE tetramer. Our cryo-EM structure reveals the basis for assembly of the physiological tetramers, and has implications for the therapeutic applications of HuBChE. This mode of tetramerization is seen only in the cholinesterases, and may provide a promising template for designing other proteins with improved circulatory residence times.

## Introduction

The cholinesterases (ChEs) are a specialised family of enzymes that hydrolyze choline-based esters, such as the neurotransmitter acetylcholine. Acetylcholinesterase (AChE) functions primarily to terminate transmission at cholinergic synapses by rapid hydrolysis of acetylcholine (1). Butyrylcholinesterase (BChE), its sister enzyme, is much more enigmatic. It is abundant in plasma and was initially believed to be an evolutionary vestige (2). It has been suggested that it functions primarily as a detoxification enzyme, thereby protecting AChE from inhibition (3, 4). It has recently been shown, however, that it inactivates the hunger hormone, ghrelin, thus reducing obesity, as well as aggression, in male mice (5, 6). Because of its ability to hydrolyze organophosphates (OPs), such as nerve agents and pesticides (3), and drugs such as cocaine (7, 8) and heroin (9–11), BChE is a promising therapeutic agent with multiple applications.

Both AChE and BChE exist in various oligomeric forms, including monomers, dimers, and tetramers (12). AChE tetramers are anchored at cholinergic synapses through association with either a collagen tail (ColQ) or a transmembrane anchor (proline-rich membrane anchor, PRiMA) (13–15). BChE exists in plasma mainly as highly glycosylated soluble tetramers (16). The use of HuBChE as a therapeutic agent for combating OP poisoning is contingent on its possessing a long circulatory half-life (17). Because its half-life depends on its ability to tetramerize, substantial effort has been invested in developing methods to produce recombinant tetrameric HuBChE (18–21).

Tetramer formation in the ChEs is mediated by assembly of four C-terminal tryptophan amphiphilic tetramerization (WAT) helices around a central proline-rich attachment domain (PRAD), which has the configuration of a polyproline II (PPII) helix and runs anti-parallel to the WAT helices (22–24). The AChE tetramerization domain itself has been crystallised in the presence of a synthetic PRAD, revealing that the WAT helices associate non-covalently with the PRAD via hydrophobic stacking and hydrogen-bonding interactions (24). In AChE, the PRAD is contributed by either ColQ in the peripheral nervous system (PNS), or by PRiMA in the central nervous system (CNS) (13–15). Mass spectrometry of HuBChE revealed the presence of polyproline-rich peptides that are 10–39 residues long. Of these, ∼70% are derived from lamellipodin, and the remaining ∼30% from up to 20 other proteins (25). Lamellipodin is a pleckstrin homology (PH) domain-containing protein that regulates the formation of actin-based cell protrusions by interacting with Enabled/Vasodilator (Ena/VASP) proteins (26, 27). Ena/VASP proteins bind their ligands through conserved proline-rich sequences, but how exactly BChE comes to be associated with lamellipodin-derived peptides remains unclear. Presumably, HuBChE is synthesized and released into the plasma as monomers, which then associate with polyproline-rich peptides to form the functional tetramer (2).

Numerous high-resolution structures are available for various forms of the catalytic subunit of AChE and BChE from several species (28). However, these structures do not provide structural information concerning the mode of assembly of functional tetramers (29, 30). Here, we use cryo-electron microscopy (cryo-EM) to determine the structure of a physiological tetrameric form of full-length, glycosylated HuBChE. This cryo-EM structure of the HuBChE tetramer provides novel and valuable insights into the molecular architecture of a biologically relevant ChE oligomer.

## Results and Discussion

To determine the structure of native, tetrameric human BChE (HuBChE), we used enzyme purified from plasma (31). HuBChE prepared in this manner is 98% tetrameric (Extended data Figure 1). A cryo-EM single-particle dataset was collected and processed as summarised in Table 1 (see Materials and Methods for details). Following 2D classification, *ab-initio* 3D reconstruction and classification were initially performed without applying symmetry. During 3D refinement, C2 symmetry was applied, resulting in the 5.7-Å reconstruction shown in Fig. 1.

**Figure 1.**
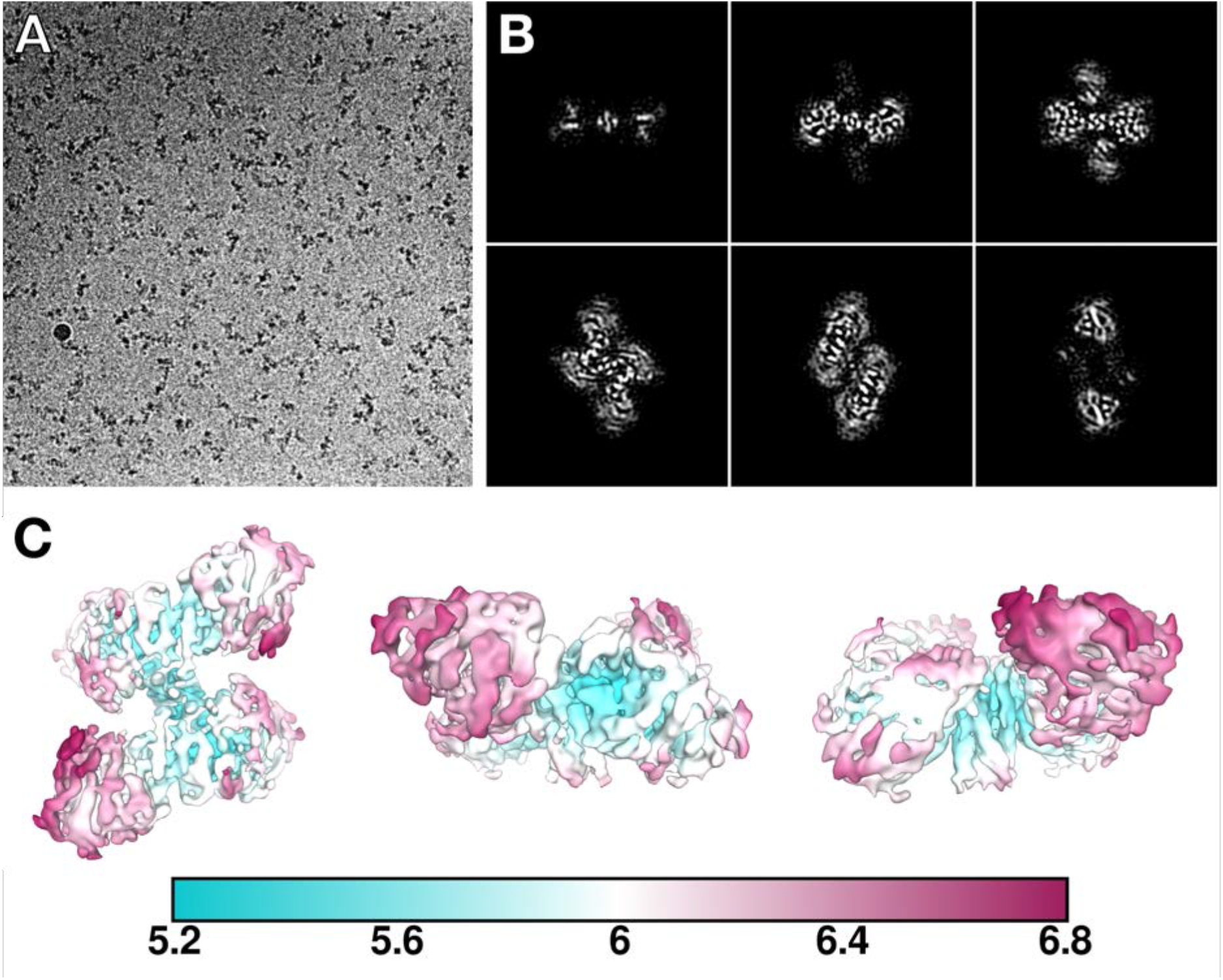
Cryo-electron microscopy reveals the organization of the HuBChE tetramer. (A) A representative micrograph from the HuBChE dataset, collected at –2.5 µm defocus and a pixel size of 1.0285 Å. (B) Slices through the 5.7-Å cryo-EM map, taken every ∼10 Å. (C) Local resolution of the HuBChE map estimated by Relion.

**Table 1.**
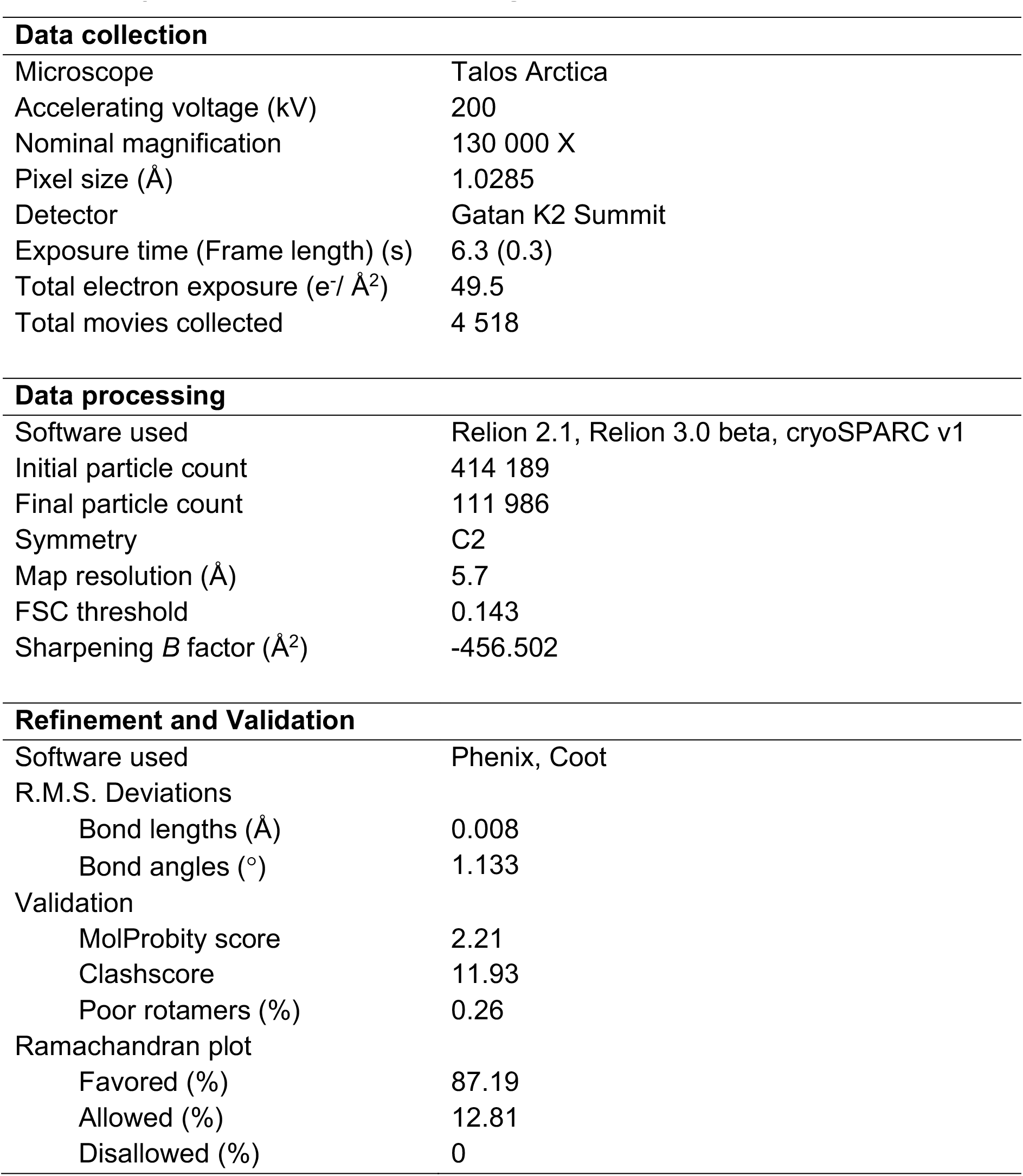
Cryo-EM Collection, Processing, Refinement, and Validation Statistics.

To model the structure of tetrameric HuBChE, four copies of the monomer, taken from the crystal structure of recombinant full-length HuBChE, PDB 2PM8 (32), were fitted as rigid bodies into the cryo-EM map. This map unambiguously shows that BChE tetramerization is mediated by assembly of the C-terminal WAT helices into a left-handed superhelix around a central polyproline II helix (Fig. 1B, Fig. 2A), strikingly similar to that of the WAT/PRAD complex of AChE (24). Consequently, a homology model could be fitted as one rigid body (see Material and Methods). Nevertheless, during real-space refinement, each of the WAT helices, as well as the PRAD, were allowed to move as independent rigid bodies in order to obtain the best fit into the cryo-EM map. Since our map is not of sufficient resolution to confidently assign side chains, the PRAD sequence was left unaltered in the current model, and thus corresponds to residues 72–86 of human ColQ (LLTPPPPPLFPPPFF), even though the primary polyproline peptide associated with HuBChE tetramers is from lamellipodin, and has the sequence PSPPLPPPPPPPPPPPPPPPPPPPPLP (25, 33). This complete HuBChE tetramer model was then global real-space refined in Phenix (34, 35), resulting in the final model shown in Fig. 2. Two-fold symmetry could not be applied during the real space refinement due to the PRAD in the centre. After refinement, the RMSD between our HuBChE WAT/PRAD and AChE WAT/PRAD (PDB 1VZJ) is 1.48 Å for Cα.

**Figure 2.**
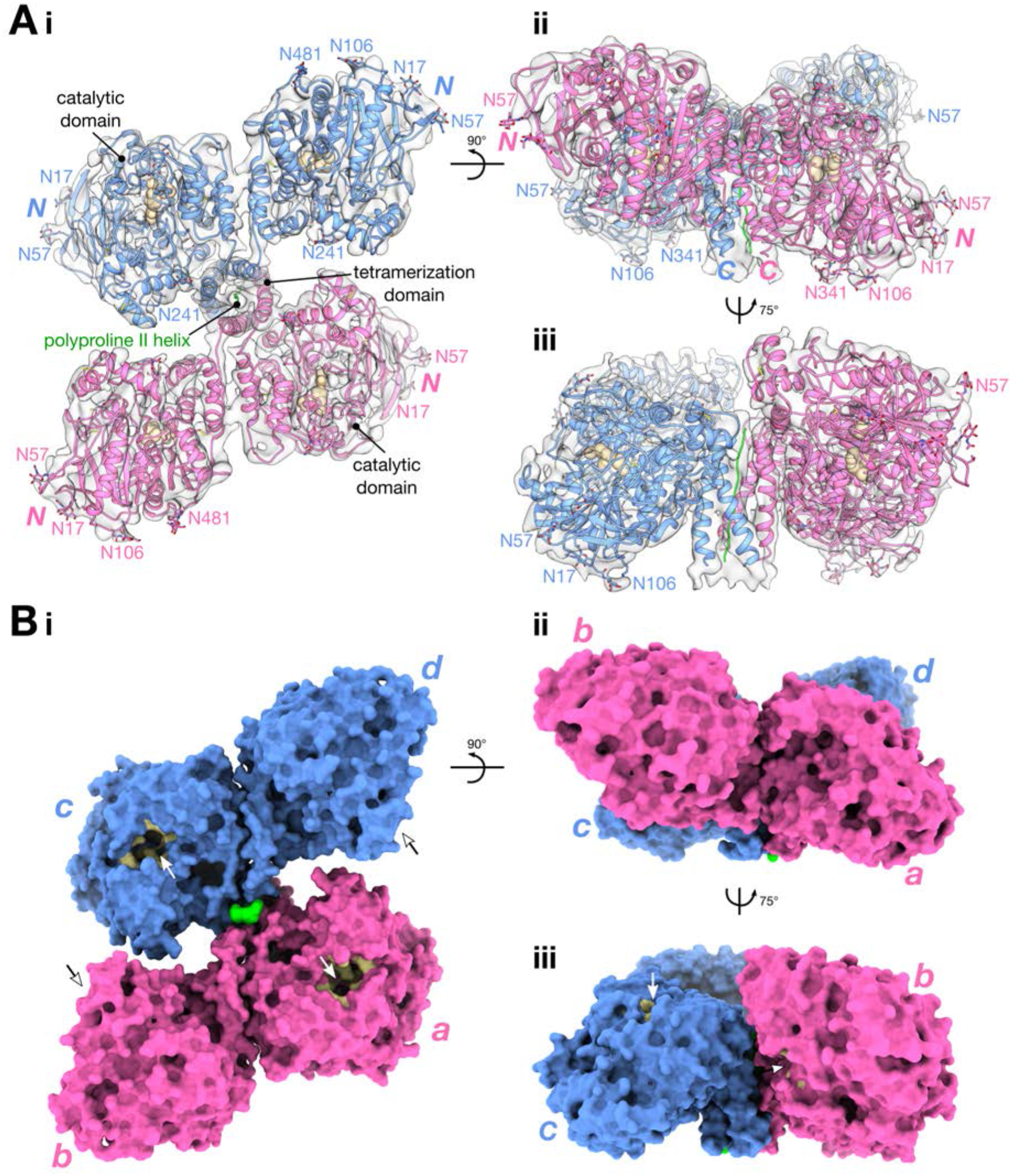
The HuBChE tetramer is a staggered dimer of dimers. The structure is presented as (A) a ribbon model fitted into the cryo-EM map (grey), and (B) a molecular surface. In (A), the catalytic triad (Ser198, Glu325, His438) is shown as tan spheres, disulfide bonds are highlighted in yellow, and glycans are shown as sticks. “N” and “C” denote the position of the N-and C-terminus of each subunit, respectively. In (B), the active-site gorge is painted in tan, and indicated by an arrow. Individual subunits are labelled *a, b, c*, and *d*, according to their chain IDs in the PDB deposition.

### The HuBChE tetramer is a staggered dimer of dimers

The HuBChE tetramer is organised as a dimer of dimers (Fig. 2). Although each monomer was fitted as a separate rigid body the resulting dimers are remarkably similar (RMSD for Cα atoms 1.13 Å) to the canonical dimer described by Brazzolotto *et al*, PDB 4AQD (36), in which individual monomers associate via a 4-helix bundle like that in the dimer observed in the prototypic *Torpedo californica* AChE (*Tc*AChE) crystal structure (37). The cryo-EM map reveals that the four catalytic domains do not lie in the same plane. Instead, each dimer is tilted ∼67° relative to the vertical axis of the WAT/PRAD PPII helix (Fig. 2Aii, Bii). This arrangement of the monomers results in a “double seesaw”-like appearance when the tetramer is viewed from the side. Furthermore, each dimer is shifted horizontally with respect to the other when viewed along the vertical axis of the tetramerization domain (Fig. 2Ai, Bi). The HuBChE tetramer thus assembles as a quasi (due to the PRAD) C2-symmetric dimer of dimers in which the monomers are diagonally equivalent.

Our structure differs from previous low-resolution crystal structures obtained for tetrameric *Ee*AChE (29, 30). These structures have RMSD values of 12.96 Å (PDB 1C2B), 13.78 Å (PDB 1EEA), and 21.84 Å (PDB 1C2O) relative to our cryo-EM model. A homology model of HuBChE constructed based on the 1C2O *Ee*AChE structure and on the 1VZJ WAT/PRAD structure (38) differs significantly from the cryo-EM structure.

The catalytic domains within a dimer are asymmetrically linked to their respective WAT helices (Fig. 3A). The break in 2-fold symmetry between the monomers occurs at G534, and is imposed by the orientation of the dimer relative to the tetramerization domain. The 4-helix bundle holding the dimer together is oriented ∼45° relative to the tetramerization domain vertical axis (Figure 3B). Consequently, for the WAT helices to form the tetramerization domain, the linkers connecting the H helices from each monomer need to run in different directions relative to their respective catalytic domains.

**Figure 3.**
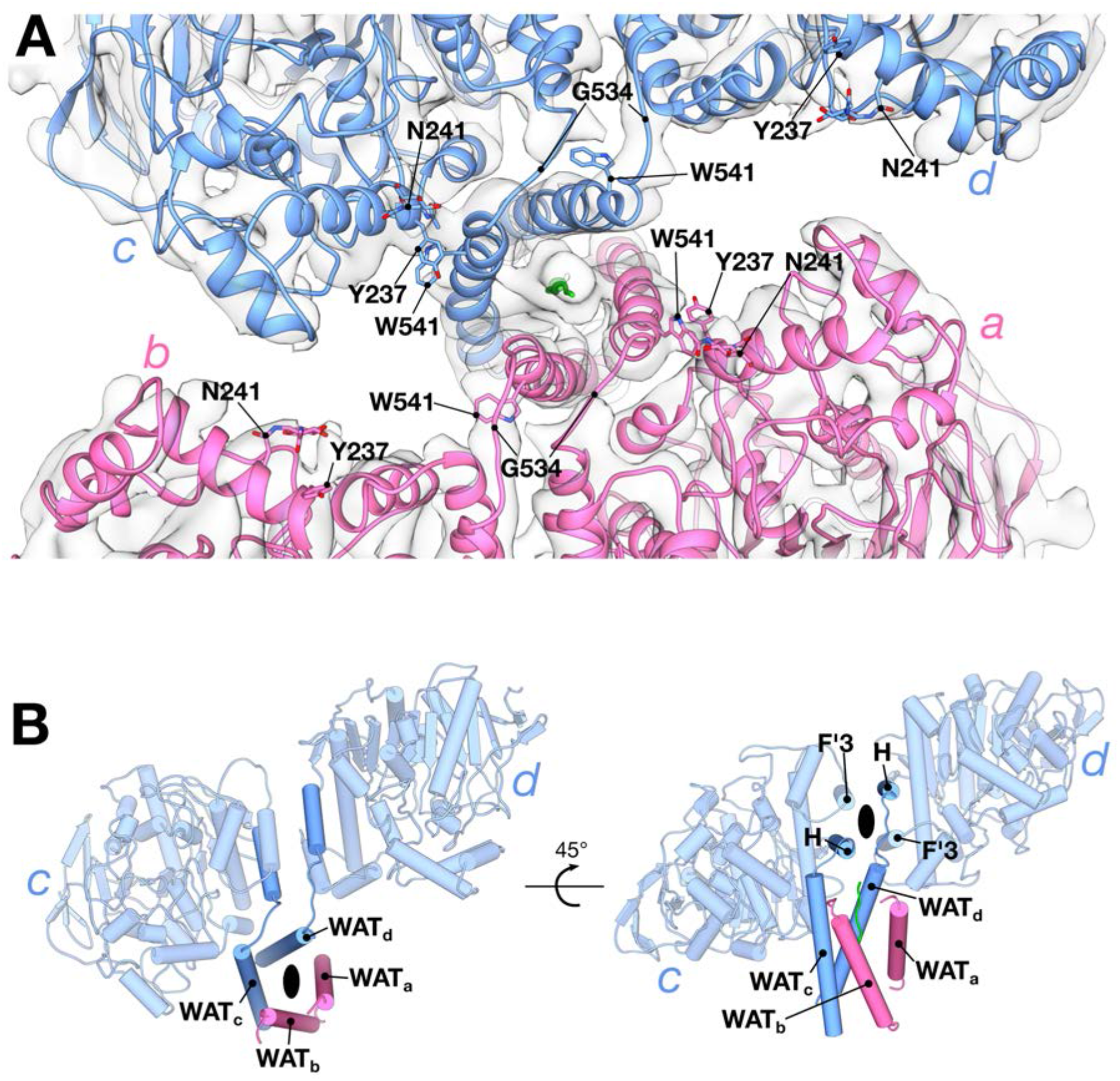
Subunits within a HuBChE dimer are asymmetrically linked to the tetramerization domain. (A) Close-up top view down the tetramerization domain. (B) Individual subunits within the dimer are asymmetrically linked to the tetramerization domain.

The ∼67° tilted assembly of the dimers around the tetramerization domain results in two different arrangements for the catalytic domains relative to the tetramerization domain (Fig. 4). In one diagonal pair of equivalent monomers (chains b and d), the catalytic domains are angled outward from the WAT helix, and thus more separated from the tetramerization domain (open arrangement; Fig. 4A), while in the other diagonal pair (chains a and c), the catalytic domains are angled inwards, towards the WAT, and are thus relatively nearer to the tetramerization domain (closed arrangement; Fig. 4B). Some of the density for the linkers connecting the catalytic domains in the closed arrangement (chains a and c) is weaker than that seen for the open arrangement (Fig. 3A).

**Figure 4.**
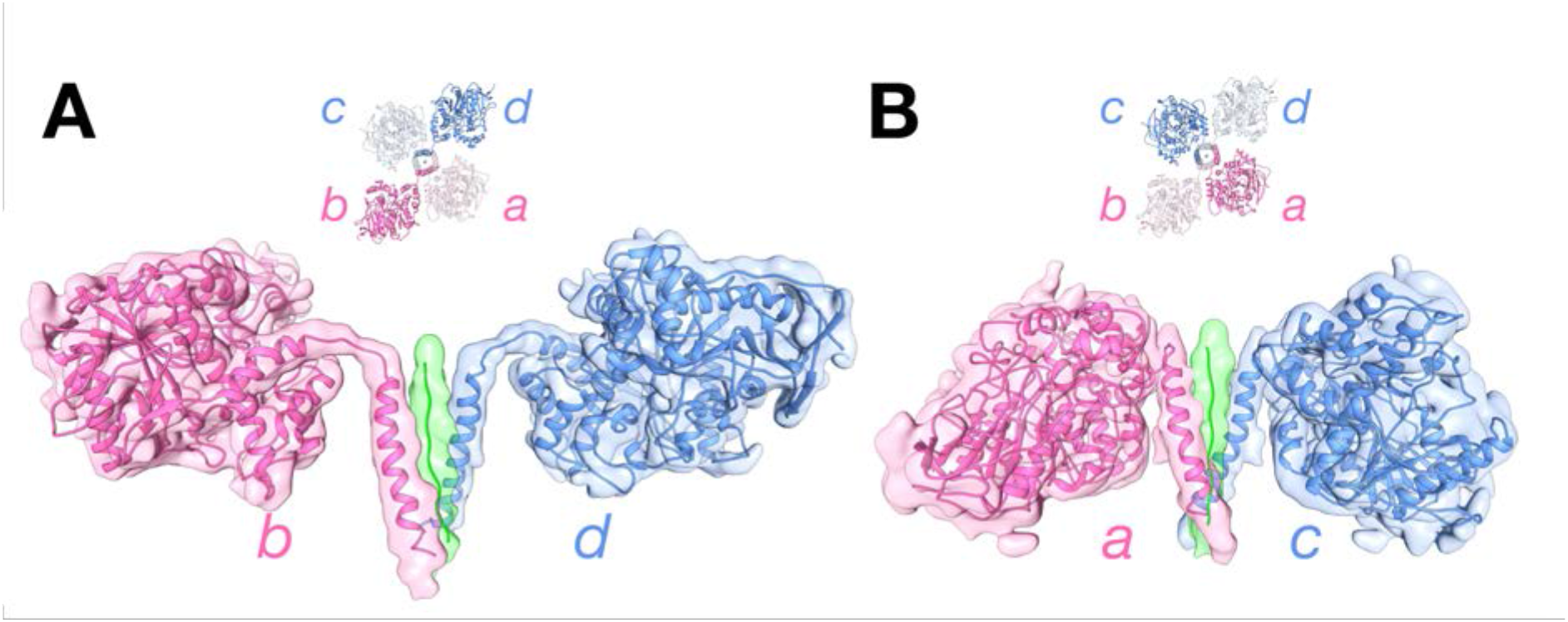
Two different arrangements for the catalytic domains relative to the tetramerization domain. Side-views of the subunits in the “open” (A) arrangement (chains b and d), and the “closed” (B) arrangement (chains a and c).

Based on local resolution estimates, the catalytic domains in the closed arrangement are slightly better-resolved than the subunits in the open arrangement (Fig. 1C). In the closed arrangement, we observed a density that can be assigned to Tyr237 of the catalytic domain and Trp541 of the WAT helix (Fig. 3A). This indicates a local interface between the catalytic domain and the tetramerization domain. It is worth noting that the equivalent residues in HuAChE are Gly242 and Arg551, respectively. Near this interface is the glycosylation site Asn241 (Fig. 3A), at which we see putative glycan density (Fig. 5E). HuAChE does not have a glycosylation site at this position, and instead has Arg246.

**Figure 5.**
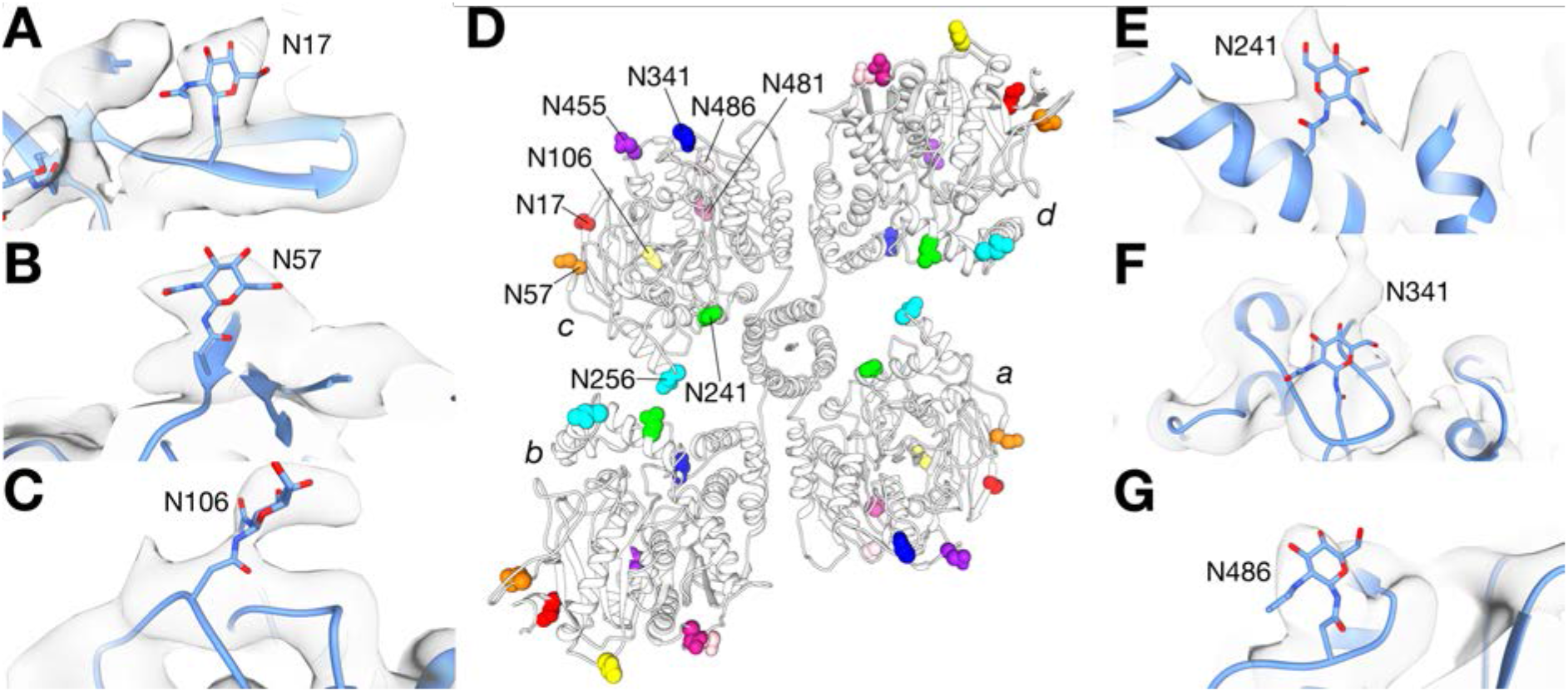
Putative glycan densities are visible in the cryo-EM map. Each HuBChE chain has nine *N*-linked glycosylation sites (43). The positions of these sites in the tetramer are shown in (D). In our map, we observe putative glycan densities at six of these sites, N17 (A), N57 (B), N106 (C), N241 (E), N341 (F), and N486 (G).

Outside the tetramerization domain, there seems to be minimal interaction between the two dimers. No surface area is buried at the dimer-dimer interface, *viz*. between the catalytic domains of chains a and d, or of chains b and c (Figure 2Bi). Solvent accessibility of the dimer-dimer interface is crucial, since the entrances to two of the active-site gorges, those of subunits b and d, are at this interface (Fig. 2Bi,iii). Indeed, the kinetic parameters of the native tetramer are close to those reported for a low-glycosylated monomeric recombinant HuBChE (39).

Overall, due to the arrangement of the catalytic subunits around the tetramerization domain, the latter does not completely protrude out of the tetramer as had been previously suggested (38). Indeed, the fact that the tetramerization domain is mostly shielded by the catalytic domains may contribute to the relative stability of the HuBChE tetramer. Thus, thermostability measurements showed that tetrameric HuBChE is more stable than tetrameric *Tc*AChE by ∼10°C (Extended Data Figure 1B,C). It should be noted that our model ends at residue 564, and lacks 10 C-terminal residues, which may partially protrude out. Importantly, these 10 residues include Cys571, which forms an interchain disulphide within the canonical dimer, and has been shown to further stabilize the tetramer (41).

We note that this mode of tetramer assembly has so far only been observed for ChEs, although it may provide a promising template on which to base the design of engineered proteins for which oligomerization, so as to extend their circulatory half-life, would be beneficial.

### The cryo-EM structure reveals differences between the HuBChE dimer and the AChE dimer

The HuBChE dimer interface is clearly formed by a 4-helix bundle comprised of helices 362–373 (helix F’3) and 516–529 (helix H) from each monomer. Alignment of the HuBChE dimer with the HuAChE dimer (PDB 3LII) (28) reveals differences in the relative orientation of subunits (Fig. 6). Although one catalytic domain of HuAChE can be superimposed well on one catalytic domain of HuBChE (RMSD = 1.56 Å) (Fig. 6A, left subunit), the second catalytic domain does not align as well (RMSD = 8.73 Å) (Fig. 6A, right subunit). This difference has implications for the relative orientations of the active-site gorges in the corresponding ChE dimers; the angle between the gorge entrances is smaller in the BChE dimer than in the AChE dimer (Fig. 5B,C).

**Figure 6.**
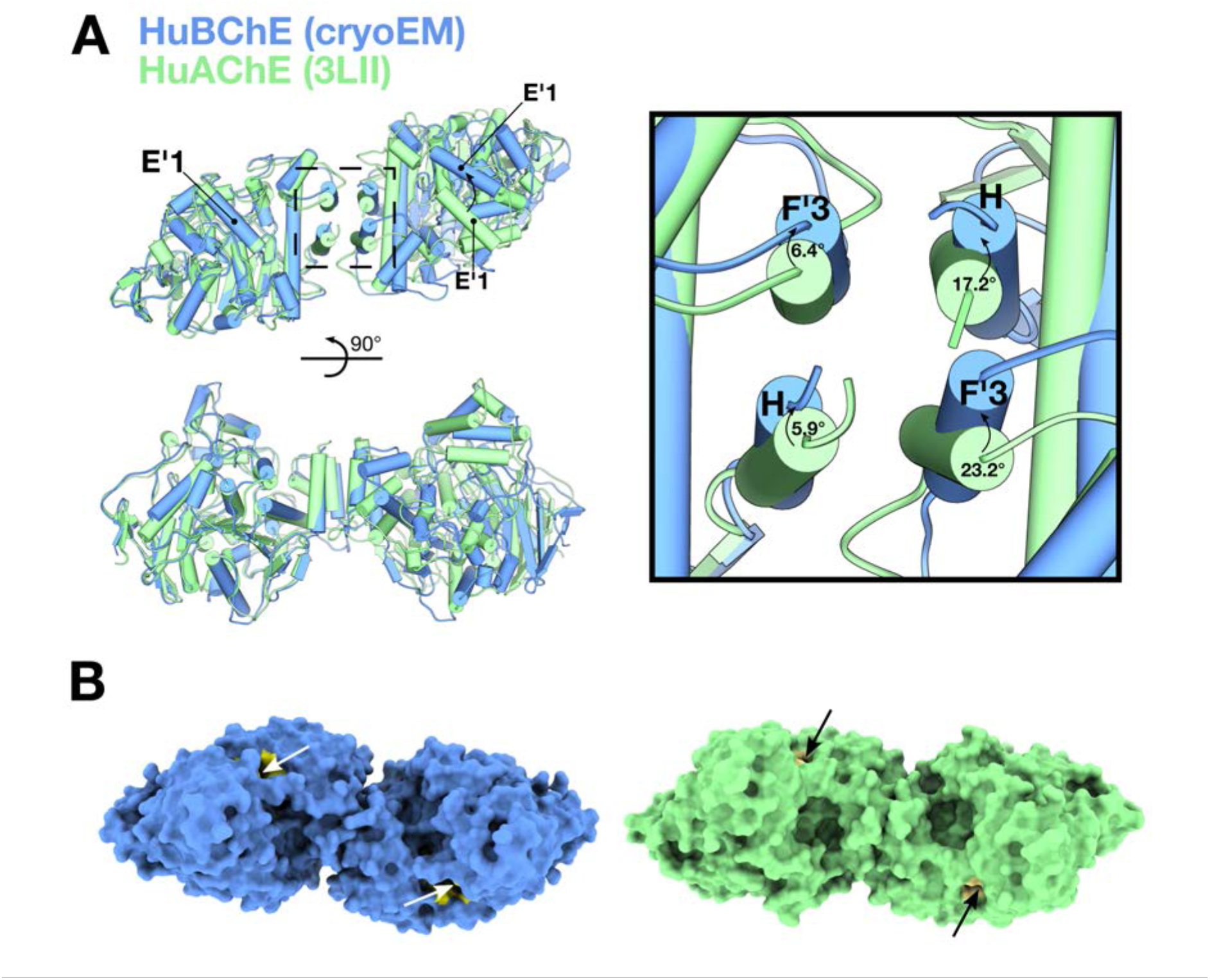
The cryo-EM structure reveals differences between BChE and AChE dimers. (A) Superposition of the HuBChE cryo-EM dimer (in blue) and the HuAChE crystal structure dimer (PDB 3LII) (in green). The inset shows the four-helix bundle at the dimer interface. Corresponding angles between the H and F’3 helixes are marked. (B) Surface representations of the HuBChE dimer and of the HuAChE dimer. In both cases, the active-site gorge is painted tan, and indicated by an arrow.

While it is possible that the differences observed between the HuBChE and HuAChE dimer structures arise simply because the structures of the AChE dimer were determined in the absence of the tetramerization domain, the structural differences between them may be functionally relevant. When AChE and BChE are co-expressed along with a PRAD peptide from PRiMA, AChE and BChE homotetramers are formed, along with “hybrid” tetramers that consist of one AChE homodimer and one BChE homodimer (42). Even though mammalian AChE can form heterodimers with avian AChE, mammalian AChE does not seem to be able to heterodimerize with mammalian BChE (42). Analysis of the residues comprising the 4-helix bundle shows that there is high sequence identity between HuBChE and HuAChE in this region. The most obvious differences are that the leucine residues at positions 373 and 380 (helix F’3) and position 536 (helix H) in HuAChE are replaced by phenylalanines at the corresponding positions 364, 371, and 526 in HuBChE.

Furthermore, there is a fundamental difference between how BChE and AChE are arranged around their respective tetramerization domains. In BChE, disulphide bonds are formed within the canonical dimers (41), and there is no covalent linkage to the proline-rich polypeptide. In AChE, there is a direct disulphide bridge between the monomers of one canonical dimer; but in the other dimer, the same residues (Cys580 of both subunits) make disulphide bonds with two conserved Cys residues adjacent to the PRAD region of ColQ and PRiMA. It would, therefore, be interesting to find out whether the architecture of the mammalian AChE tetramer is similar to that of the BChE tetramer presented here.

## Concluding Remarks

We used cryo-EM to determine the structure of a physiological tetrameric form of a ChE, that of HuBChE. Our experimental structure clearly shows that the tetramer is a markedly non-planar staggered dimer of dimers, with tetramerization mediated by assembly of the C-terminal WAT helices around a central polyproline II helix. Our findings raise questions concerning differences in oligomeric interactions within AChE and BChE. In particular, the cryo-EM HuBChE tetramer structure differs from previous models based on low-resolution crystal structures of tetrameric *Electrophorus* AChE (29, 30, 38), which placed the four monomers in roughly the same plane, and thus differ significantly from the cryo-EM structure. It is thus important that cryo-EM structures should be determined for the G_4_ and collagen-tailed (A_12_) forms of AChE. Use of an oligomeric polyproline II-like helix around which amphiphilic a-helical sequences wrap as a super-helical motif could be used as a generic tool to stabilize other proteins as tetramers, with the potential to extend their pharmacokinetic lifetimes.

## Materials and Methods

### Purification of HuBChE from Plasma

Outdated human plasma from the University of Nebraska Medical Center blood bank was purified by anion exchange chromatography at pH 4.0, followed by affinity chromatography on procainamide-Sepharose at pH 7.0 (31), and affinity chromatography on a Hupresin column at pH 8.0. UniProt P06276 provides the amino acid sequence of 28 residues in the signal peptide and 574 residues in the mature, secreted 85-kDa BChE subunit (43). The HuBChE protein was 99% pure, and consisted predominantly of tetramers, as visualized on a non-denaturing gel stained for BChE activity according to Karnovsky and Roots (44) (Extended Data Figure 1).

### Thermal Unfolding Assays

Thermostability of HuBChE was assessed by nano-differential scanning fluorimetry (nanoDSF), performed on a Prometheus NT.48 (NanoTemper) using a temperature ramp of +3°C/min. Approximately 10 µL of purified protein was loaded into nanoDSF-grade high sensitivity capillaries (NanoTemper). HuBChE was compared to purified G_4_ *Torpedo californica* AChE (*Tc*AChE). HuBChE samples were measured in 10 mM Tris, pH 8.0, at 3.3 mg/mL, and *Tc*AChE samples in 1 M NaCl, 10 mM Tris, pH 8.0, at 2 mg/mL.

### Cryo-EM Grid Preparation

HuBChE, at 3.3 mg/mL in 10 mM Tris, pH 8.0, was used for cryo-EM. Approximately 3 µL protein solution was pipetted onto glow-discharged 300-mesh Quantifoil R1.2/1.3 holey carbon grids, which were blotted for either 3 s at blot force 0, or 2 s at blot force –2, on a Vitrobot Mark IV (FEI) at 20°C and 95% humidity. Grids were immediately plunged into liquid ethane cooled to liquid N_2_ temperature, and stored under liquid N_2_ prior to imaging.

### Cryo-EM Data Collection

Data collection was performed on a Talos Arctica (FEI) transmission electron microscope operated at 200 kV, equipped with a postcolumn energy filter (Gatan) operated in zero-loss imaging mode with a 20-eV energy-selecting slit. All images were recorded on a postfilter K2 Summit direct electron detector (Gatan). Movies were collected using EPU (FEI) in super-resolution counting mode, at a nominal magnification of 130 000 X, corresponding to a pixel size of 1.0285 Å at the specimen level. The exposure time for each movie was 6.3 s with 0.3-s frames. The dose per frame was 2.36 e-/Å^2^, resulting in a total dose of 49.5 e-/Å^2^. A total of 4,518 movies were collected, with defocus targets ranging from –1.2 to –2.5 µm.

### Cryo-EM Image Processing

A total of 4,518 movies were imported into Relion 2.1.0 (45). Drift and beam-induced motion were corrected with MotionCor2 (46). Contrast transfer function (CTF) estimation was performed with GCTF 1.06 (47). All micrographs with a reported maximum resolution worse than 8 Å were discarded, resulting in a set of 3,267 micrographs. From these, 832 particles were manually selected, and used to generate 2D classes, which were then used as references for automated particle picking. A total of 414,189 particles were picked and subjected to two rounds of 2D classification. After 2D classification, 374,086 particles were selected and imported into cryoSPARC (48) for *ab-initio* 3D classification without applying symmetry. The class showing the highest resolution was selected, and the particles were refined with C2 symmetry. The resulting map was then low-pass filtered to 50 Å, and used as a template for 3D classification in Relion 3-beta. After two rounds of 3D classification, 111,986 particles were selected for auto-refinement in Relion 3-beta, using C2 symmetry. Refinement was performed with a soft mask prepared by low-pass filtering the cryoSPARC map to 15 Å, and adding a 10-pixel extension and a 10-pixel soft cosine edge. After postprocessing and sharpening with a *B*-factor of –456.502, the resulting map had a resolution of 5.7 Å (Extended Data Figure 2), based on the “gold standard” FSC = 0.143 criterion (49). Local resolution was estimated in Relion 3-beta.

### Model Building

Rigid body fitting was performed in Chimera, followed by refinement in Phenix (34, 35). Four HuBChE monomers from PDB 2PM8 were fitted into the cryo-EM map. In order to generate a starting model of full-length HuBChE to use in later model building, homology models of the WAT helices, comprising residues 535–574, were constructed with MODELLER (50), using the WAT domain of AChE (PDB 1VZJ) as a template. The homology model of the tetramerization domain was first fitted as one rigid body. During the first round of real space refinement the model was rigid body refined, resulting in 9 independent rigid bodies (4 catalytic domains, 4 WAT helices, and 1 PRAD). Residue 534 of the catalytic domain and residue 535 of the WAT helix were connected and real-space refined with Coot (51). Residues Q17, Q455, Q481, and Q486 in the starting models of the catalytic domains (from PDB 2PM8) were restored to N residues, as they are in wild-type HuBChE. Glycans were as in PDB 4AQD. The complete BChE model was then globally real-space refined in Phenix (34, 35), with secondary structure restraints but without non-crystallographic symmetry (NCS) restraints that could not be applied due to the PRAD. Figures were prepared with Chimera (52) and Chimera X (53).

## Acknowledgements

The authors sincerely acknowledge Dr. M. Vanevic for IT support, and Engr. C.W.T.M. Schneijdenberg, J.D. Meeldijk, and Dr. S.C. Howes for management and maintenance of the Utrecht University EM-Square Facility.

---

**Extended Data Figure 1.**
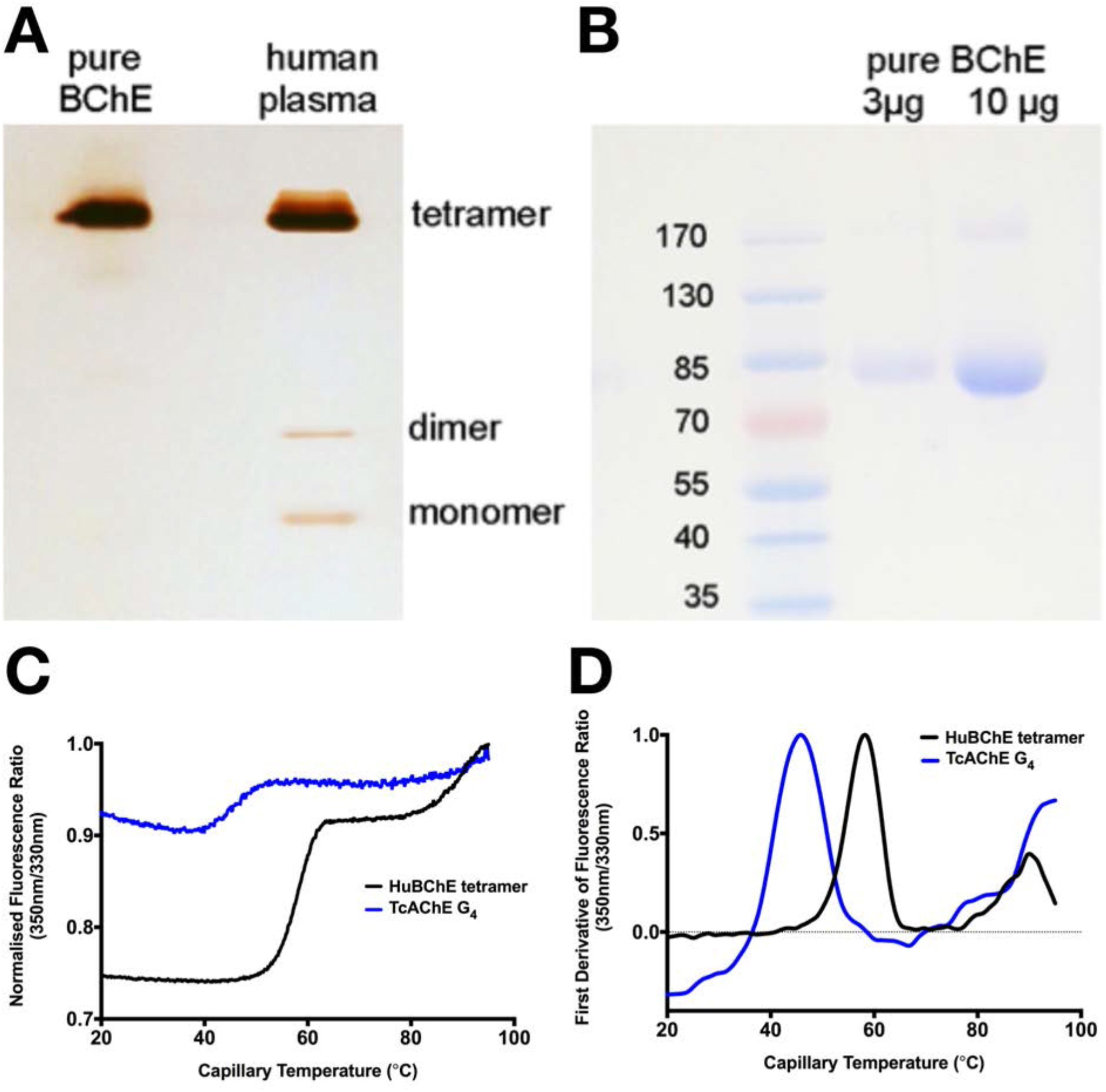
Purity and sample quality of HuBChE tetramers used for cryo-EM. (A) A non-denaturing polyacrylamide gel stained for BChE activity with butyrylthiocholine, according to Karnovsky and Roots (44). The band of BChE activity in pure HuBChE migrates to the same position as the dominant band in human plasma, which is known to be the HuBChE tetramer (26). (B) SDS-polyacrylamide gel stained with Coomassie blue shows a band at 85 kDa for the HuBChE monomer produced by boiling the tetramer in 2% SDS/50 mM dithiothreitol, and a weak band at 170 kDa for the non-reducible dimer. (C,D) NanoDSF thermostability assays of purified HuBChE and G_4_ *Tc*AChE. Results are shown as (C) the ratio of fluorescence intensity at 350 nm to that at 330 nm, and as (D) the first derivative of this ratio.

**Extended Data Figure 2.**
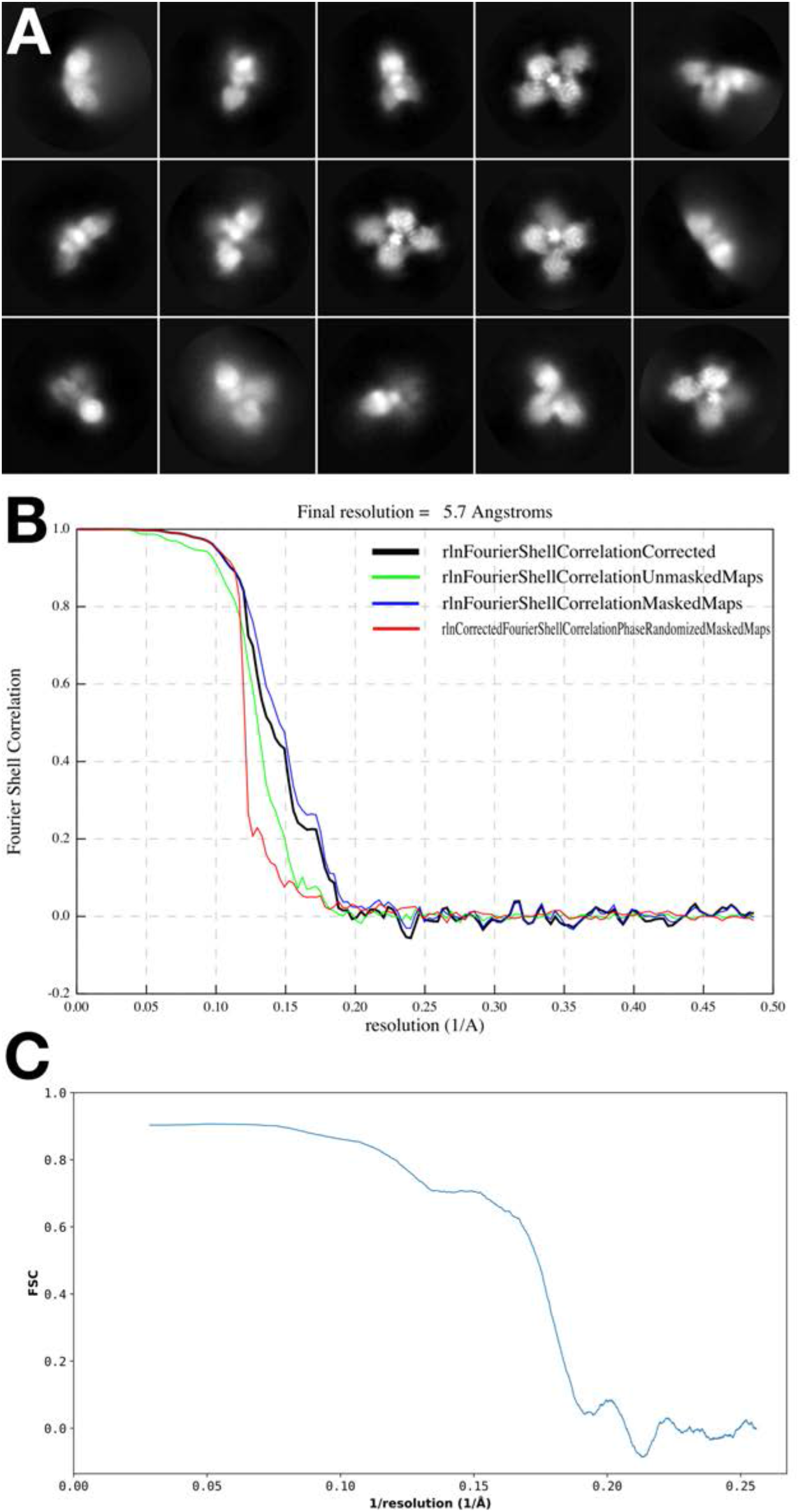
Representative 2D class averages and resolution estimates of the HuBChE map. (A) Representative 2D class averages from the HuBChE dataset. (B) FSC curves from PostProcessing in Relion 3-beta. (C) Map vs model FSC curves from Phenix.

